# Transport Phenomena in Fluid Films with Curvature Elasticity

**DOI:** 10.1101/2020.01.14.906917

**Authors:** Arijit Mahapatra, David Saintillan, Padmini Rangamani

## Abstract

Cellular membranes are elastic lipid bilayers that contain a variety of proteins, including ion channels, receptors, and scaffolding proteins. These proteins are known to diffuse in the plane of the membrane and to influence the bending of the membrane. Experiments have shown that lipid flow in the plane of the membrane is closely coupled with the diffusion of proteins. Thus there is a need for a comprehensive framework that accounts for the interplay between these processes. Here, we present a theory for the coupled in-plane viscous flow of lipids, diffusion of transmembrane proteins, and curvature elastic deformation of lipid bilayers. The proteins in the membrane are modeled such that they influence membrane bending by inducing a spontaneous curvature. We formulate the free energy of the membrane with a Helfrich-like curvature elastic energy density function modified to account for the chemical potential energy of proteins. We derive the conservation laws and equations of motion for this system. Finally, we present results from dimensional analysis and numerical simulations and demonstrate the effect of coupled transport processes in governing the dynamics of membrane bending and protein diffusion.

## 1 Introduction

Lipid bilayers are present both in the plasma membrane and in intracellular organelles [1] and have an extremely heterogeneous composition [2]. They consist of many different types of lipids, integral, and peripheral membrane proteins [3], all of which are important in cellular function [4]. One of the classic features of cellular membranes is their ability to bend out of plane and this has been the focus of many studies, both theoretical and experimental, over the past five decades [5, 6, 7, 8, 9]. We now also know that these membrane-protein interactions in cells are associated with many curvature sensing [10] and curvature generating phenomena [11] including tubulation [12], vesicle generation [13], and membrane trafficking [14, 15]. Proteins embedded in the membrane diffuse in the plane of the membrane and undergo transport by advection processes associated with lipid flow [16]. Experimental observations in reconstituted or synthetic lipid vesicles show that the coupling of lipid flow, protein diffusion, and membrane bending can give rise to emergent phenomena [17, 18, 19]. Thus, there is a need to understand how protein diffusion, lipid flow, and membrane bending are coupled to determine the mechanical response of lipid bilayers.

Many groups have focused on the development of theoretical models of lipid bilayer mechanics [20, 21, 22]. The seminal work of [23], [24], and [25] established the framework for using variational principles and thin shell mechanics for modeling membrane bending. Later, [26] established the correspondence between Koiter’s shell theory and developed a complete theoretical framework of membrane mechanics. These early models assumed the membrane to be inviscid and focused primarily on the elastic effects. In the past decade, many groups have proposed the addition of viscous effects in addition to membrane bending [27, 28, 29, 30] building on the ideas proposed by [31]. We also showed recently that including intrasurface viscosity in addition to membrane bending allows for the calculation of local membrane tension in the presence of protein-induced spontaneous curvature [32] and for the calculation of flow fields on minimal surfaces [33]. Separately, the interaction between in-plane protein diffusion and membrane bending has been modeled [34, 35, 36, 37, 38, 13]. Specifically, [39] proposed a framework that included the chemical potential energy of membrane-protein interactions and membrane bending and demonstrated the interaction between bending and diffusion. A series of studies by Arroyo and coworkers also developed a comprehensive framework for incorporating membrane-protein interactions using Onsager’s variational principles [40, 30, 41].

Building on these efforts, we provide a coupled theory of both in-plane viscous flows and diffusion of curvature-inducing transmembrane proteins coupled with membrane bending. We note that a version of this model was presented in [42]. Using a free energy functional that includes bending energy, chemical potential energy of membrane-protein interactions, and intrasurface viscosity, we derive the governing equations of motion in § 2. In § 3, we analyze this system of equations assuming small deformations from the flat plane and identify the role of different dimensionless groups in governing the regimes of operation. We then perform numerical simulations in a one-dimensional model in § 4.1 and in a two-dimensional Monge parametrization in § 4.2. These results are cast in perspective of the current knowledge of the field and future directions are presented in § 5.

## 2 Membranes with intra-surface viscosity and protein diffusion

We formulate the governing equations for the dynamics of an elastic lipid membrane with surface flow, coupled to the transport of membrane-embedded proteins that induce spontaneous mean curvature. The notations used in the model are summarized in Table 1. We assume familiarity with tensor analysis and curvilinear coordinate systems [43, 44, 45].

**Table 1:** Summary of the notations used in the model.

| Notation | Description | Units |
| --- | --- | --- |
| $\gamma$ | Lagrange multiplier for incompressibility | $\text{pN} \cdot \text{nm}^{-1}$ |
| $p$ | Pressure difference across the membrane | $\text{pN} \cdot \text{nm}^{-2}$ |
| $C$ | Protein-induced spontaneous curvature | $\text{nm}^{-1}$ |
| $\theta^\alpha$ | Surface coordinates | |
| $W$ | Local free energy per unit area | $\text{pN} \cdot \text{nm}^{-1}$ |
| $\lambda$ | Membrane tension, $\lambda = -(W + \gamma)$ | $\text{pN} \cdot \text{nm}^{-1}$ |
| $\lambda_0$ | Membrane tension at infinity | $\text{pN} \cdot \text{nm}^{-1}$ |
| $\zeta^{\alpha\beta}$ | Elastic stress tensor | $\text{pN} \cdot \text{nm}^{-1}$ |
| $\pi^{\alpha\beta}$ | Viscous deviatoric stress | $\text{pN} \cdot \text{nm}^{-1}$ |
| $H$ | Mean curvature of the membrane | $\text{nm}^{-1}$ |
| $K$ | Gaussian curvature of the membrane | $\text{nm}^{-2}$ |
| $k$ | Bending modulus (rigidity) | $\text{pN} \cdot \text{nm}$ |
| $\bar{k}$ | Gaussian modulus | $\text{pN} \cdot \text{nm}$ |
| $\sigma$ | Protein density per unit area | $\text{nm}^{-2}$ |
| $\sigma_s$ | Saturation protein density per unit area | $\text{nm}^{-2}$ |
| $\ell$ | Proportionality constant of $C - \sigma$ relation | $\text{nm}$ |
| $D$ | Protein diffusion coefficient, $D = k_B T / f$ | $\text{nm}^2 \cdot \text{s}^{-1}$ |
| $f$ | Hydrodynamic drag coefficient of a protein | $\text{pN} \cdot \text{s} \cdot \text{nm}^{-1}$ |
| $L$ | Size of the domain | $\text{nm}$ |
| $k_B$ | Boltzmann constant | $\text{pN} \cdot \text{nm} \cdot \text{K}^{-1}$ |
| $T$ | Temperature | $\text{K}$ |

### 2.1 Membrane geometry, kinematics and incompressibility

The lipid membrane is idealized as a two-dimensional manifold Ω in three-dimensional space. Material points on Ω are parametrized by a position field ***r***(*θ^α^, t*), where *θ^α^* are surface coordinates and play a role analogous to that of a fixed coordinate system used to parametrize a control volume in the Eulerian description of classical fluid mechanics. Here and henceforth, Greek indices range over {1, 2} and, if repeated, are summed over that range. The local tangent basis on the surface is naturally obtained as ***a**_α_* = ***r**,_α_* where commas identify partial derivatives with respect to *θ^α^*. The unit normal field is then given by ***n** = **a***_1_ × ***a***_2_/|***a***_1_ × ***a***_2_|. The tangent basis also defines the surface metric *a_αβ_* = ***a**_α_ **· a**_β_* (or coefficients of the first fundamental form), a positive definite matrix, which is one of the two basic variables in surface theory. The other is the curvature *b_αβ_* (or coefficients of the second fundamental form) defined as *b_αβ_* = ***n · r**,_αβ_*. Of special interest are the mean and Gaussian curvatures, which will enter the Helfrich energy of the membrane and are defined, respectively, as

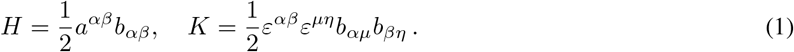

Here, *a^αβ^* = (*a_αβ_*)^−1^ is the dual metric, and *ε^αβ^* is the permutation tensor defined as 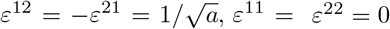.

We assume that the surface Ω is moving with time, and the velocity of a material point in the membrane is given by 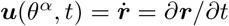. It can be expressed in components on the natural basis introduced above:

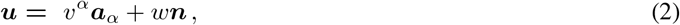

where the components *υ^α^* capture the tangential lipid flow and *w* is the normal surface velocity. The membrane is assumed to be incompressible, which prescribes a relationship between the in-plane velocity field and the curvature as [27, 29]

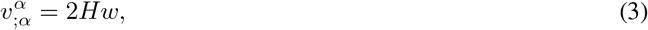

where the semi-colon refers to covariant differentiation with respect to the metric *a_αβ_*.

### 2.2 Stress balance and equations of motion

We model the membrane as a thin elastic shell and, in the absence of inertia, the equations of motion are the equations of mechanical equilibrium. For a membrane subjected to a lateral pressure difference *p* in the direction of the unit normal ***n***, these may be summarized as [26]

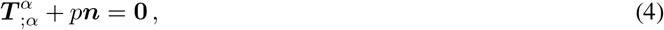

where ***T**^α^* are the so-called *stress vectors*. The differential operation in equation (4) is the surface divergence

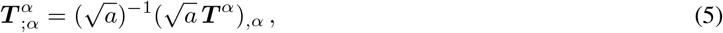

where *a* = det(*a_αβ_*) > 0. This framework encompasses all elastic surfaces for which the energy density responds to metric and curvature. For example, if the energy density per unit mass of the surface is *F*(*a_αβ_, b_αβ_*), then [26]

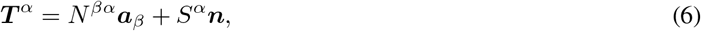

where

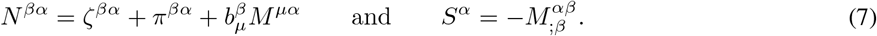

We have introduced the notation 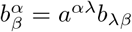. In equation (7), *ζ^βα^* is the in-plane elastic stress tensor, *π^βα^* is the intra-membrane viscous stress tensor due to surface flow, and *M^αβ^* is the moment tensor due to curvature-induced elastic bending. We discuss constitutive equations for these various contributions in the next sections.

Substituting equations (6) and (7) into (4), invoking the Gauss and Weingarten equations ***a**_β;α_* = *b_βα_**n*** and 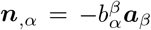 [43], and projecting the result onto the tangent and normal spaces of Ω provides the three governing equations

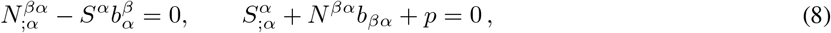

which express stress balances in the tangential and normal directions.

### 2.3 Free energy of an elastic membrane with curvature-inducing proteins

The elastic contribution of the surface stress and the moment tensor are derived from a free energy and are expressed as [26]

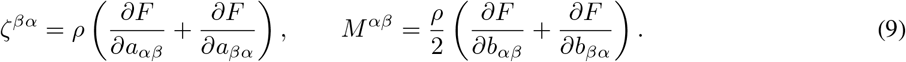

Here, *F* is the energy Lagrangian per unit mass defined as [26]

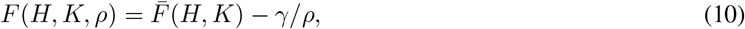

where *γ* is a Lagrange multiplier imposing the constraint of incompressibility, and *ρ* is the membrane density which is assumed to be constant. It is customary to formulate the mechanics in terms of the free energy per unit area as [46]

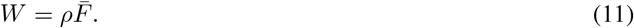

For an elastic membrane with a density *σ* of curvature-inducing proteins, we model this free energy as the sum of elastic and chemical energies [36, 39, 22]

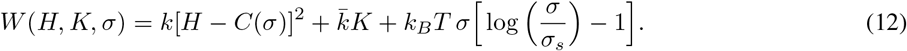

The first two terms correspond to the classical Helfrich free energy and involve the two bending moduli *k* and 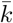. While these could in general depend on *σ*, we take them to be constant as is appropriate in the dilute limit. *C*(*σ*) is the protein-induced spontaneous curvature and is assumed to depend linearly on protein density [36, 39, 47]:

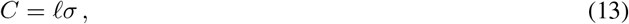

where the constant *ℓ* is a characteristic length scale associated with the embedded protein. The last term in equation (12) is the entropic contribution due to thermal diffusion of proteins [47], where *k_B_T* is the thermal energy and *σ_s_* denotes the saturation density of proteins on the membrane.

Inserting equation (12) for the free energy into equation (9) provides expressions for the elastic stress and moment tensors as

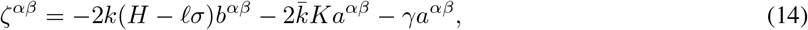

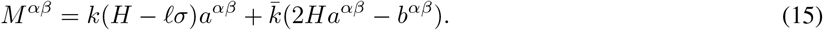

### 2.4 Viscous stress

In the presence in surface flow, a viscous stress also develops in the membrane. The deviatoric part of the viscous stress tensor *π^αβ^* is assumed to depend linearly on strain rate as [31]

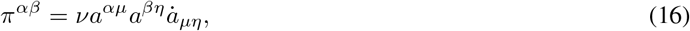

where *ν*, a positive constant, is the intra-membrane surface viscosity. Further expanding this expression, we obtain

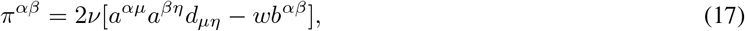

where *d_μη_* = (*υ_μ;η_* + *υ_η;μ_*)/2 is the surface rate-of-strain tensor.

### 2.5 Conservation equation for protein transport

To complete the model, we specify a transport equation for the protein density *σ*. Mass conservation can be expressed as

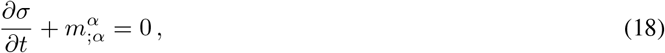

where *m^α^* denotes the protein flux. This flux has contributions from advection by the surface flow as well as from gradients in chemical potential. Following [39], it can be derived from first principles as

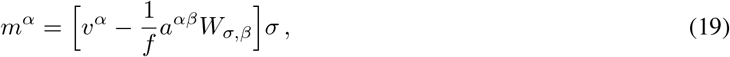

where *f* is the hydrodynamic drag coefficient of a protein and *W_σ_* = *∂W/∂σ*.

### 2.6 Summary of the governing equations

We summarize the governing equations for the membrane and protein dynamics. The tangential momentum balance, obtained by inserting equations (14), (15) and (17) for the stresses into the first equation in (8), is expressed as

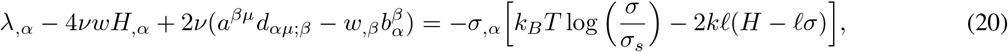

where we have introduced the membrane tension λ = −(*W* + *γ*). Along with the surface incompressibility condition

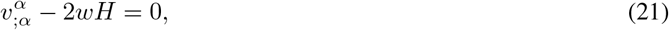

equation (20) constitutes the governing equation for the intra-membrane flow. We note the similarity with the Stokes equations, where the tension λ plays a role analogous to the pressure in classical incompressible flow [33]. The right-hand side captures the forcing by the protein distribution on the flow. Similarly, the normal force balance in (8) provides the so-called *shape equation*, written after simplifications as

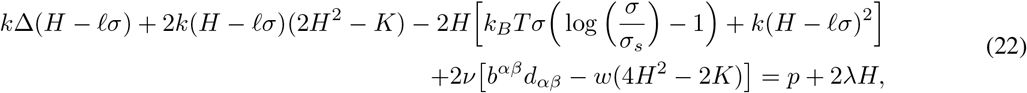

where Δ(·) = (·)*_;αβ_a^αβ^* is the surface Laplacian. Equation (22) can be interpreted as the governing equation for the position field ***r***. Finally, the model is completed by the advection-diffusion equation for the protein density, which is written

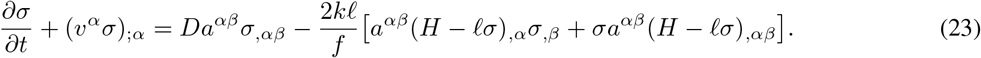

The first term on the right-hand side captures Fickian diffusion of proteins, with diffusivity *D* given by the Stokes-Einstein relation: *D = k_B_ T/f*. The second term captures the interaction of the curvature gradient and the protein gradient. The third term captures the effect of membrane shape on protein transport: mismatch between the mean curvature and the protein-induced spontaneous curvature serves as a source term for the transport of protein on the membrane surface.

## 3 Linearization and dimensional analysis

In this section, we specialize the governing equations presented in § 2.6 in a Monge parametrization assuming small deflections from the flat plane.

### 3.1 Governing equations in the linear deformation regime

The surface parametrization for a Monge patch is given by

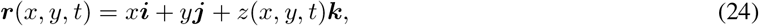

where unit vectors (***i, j, k***) form a fixed Cartesian orthonormal basis, and *z*(*x, y, t*) is the deflection from the (*x, y*) plane. The tangent and normal vectors are given by

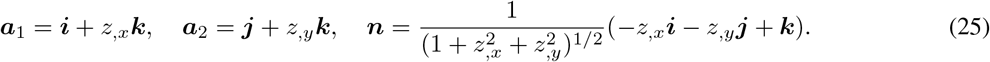

The surface metric (*a_αβ_*) and curvature metric (*b_αβ_*) take the following forms

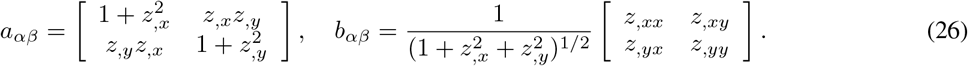

We further assume that deflections of the membrane from the flat configuration are small and simplify the governing equations in the limit of weak surface gradients |**∇**_*z*_| ≪ 1 by neglecting quadratic terms in |**∇**_*z*_| [48]. In this limit, differential operators in the space of the membrane reduce to the Cartesian gradient, divergence and Laplacian in the (*x, y*) plane. The linearized governing equations for the intra-membrane flow become:

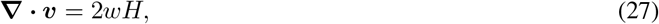

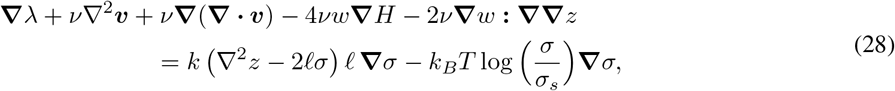

whereas the shape equation expressing the normal momentum balance is

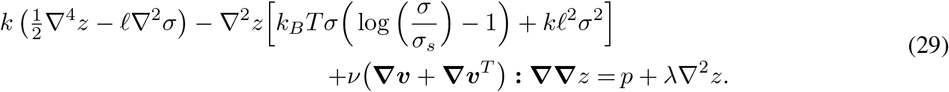

The transport equation for the protein density simplifies to

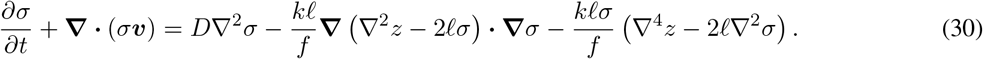

We also note the linearized kinematic relation for the normal velocity:

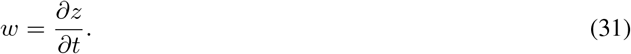

### 3.2 Non-dimensionalization

We scale this system of equations using the following reference values. Length is non-dimensionalized by the size *L* of the domain, protein density by its reference value *σ*_0_, and membrane tension by its far-field value λ_0_. We also use the characteristic velocity scale *υ_c_* = λ_0_*L/ν* and time scale *t_c_* = *L*^2^/*D*. Denoting dimensionless variables with a tilde, the scaled governing equations are:

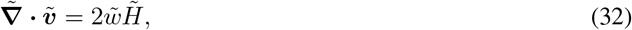

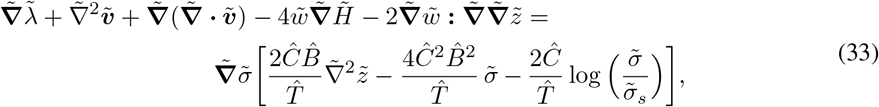

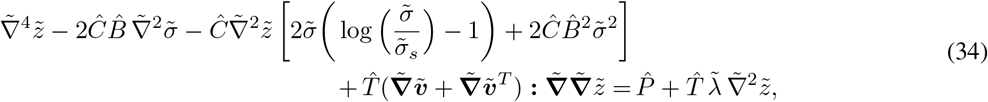

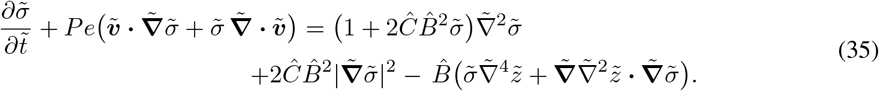

The expression for the normal velocity also becomes:

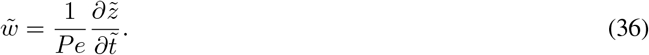

The dynamics are governed by five dimensionless parameters defined as follows. The ratio of the chemical potential to the bending rigidity of the membrane is denoted by 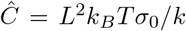. The ratio of the length scale induced by the proteins and the membrane domain is given by 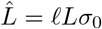. The ratio of the intrinsic length scale of the membrane to the domain size is given by 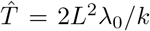. The ratio between the bulk pressure and bending rigidity is denoted by 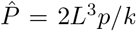. Finally, the Péclet number *Pe* = λ_0_*L*^2^/*νD* compares the advective transport rate to the diffusive transport rate. We define 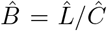 for convenience of simulations and cast the equations in terms of 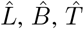, and *Pe*. Further, we assume that there is no pressure difference across the membrane 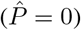.

## 4 Numerical simulations

We illustrate the interplay between protein diffusion and membrane bending using 1D and 2D numerical simulations.

### 4.1 One-dimensional simulations

We first explore the interplay between membrane bending and protein diffusion in the special case of a membrane that deforms as a string in one dimension, with a shape parameterized as 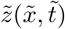. The flow of lipids does not play a role in this scenario, and as a result in-plane velocity-dependent terms vanish in equations (32)–(35). The system of governing equations reduces to

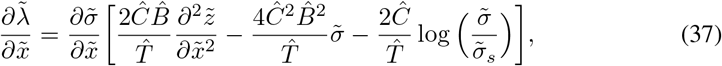

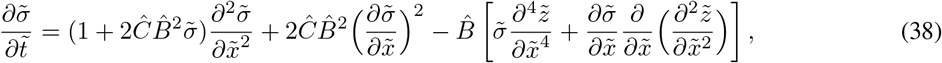

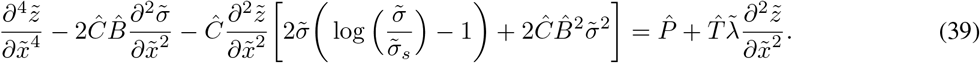

Equations (37)–(39) are solved numerically using a finite-difference scheme coded in Fortran 90. The tangential momentum balance (37), which can be viewed as an equation for the tension 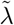, is solved subject to the condition 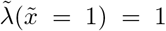, whereas the shape equation (39) is solved subject to clamped boundary conditions 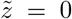 and 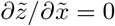 at both ends of the domain 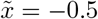, 0.5.

We first analyzed the evolution of a symmetric patch of protein defined as 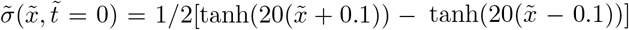, subject to no-flux boundary conditions on 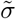 at the ends of the domain. Results from these simulations are shown in figure 1. In response to this protein distribution, the initial configuration of the membrane is bent (see figure 1(*a*) at *t* = 0). Over time, 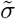 homogenizes as a result of diffusion, and therefore the deflection 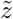 decreases. At steady state, the distribution of protein is uniform on the membrane and Z is everywhere zero. The time evolution of 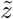 at the center of the string, corresponding to the maximum deflection, is shown in figure 1(*b*).

**Figure 1:**
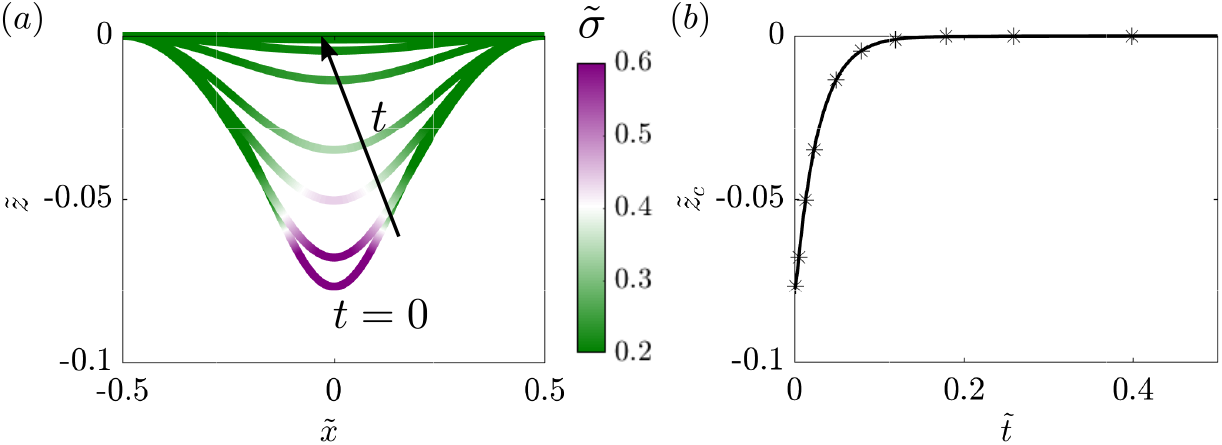
Protein density and membrane deformation in one dimension as functions of time, when an initial protein distribution and no-flux boundary conditions are prescribed. (*a*) Distribution of protein density plotted on the deformed one-dimensional membrane. (*b*) Time evolution of the maximum deflection of string 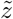.

As a second example, we discuss the case where the protein density is initially zero and a time-dependent protein flux is prescribed at both boundaries as shown in figure 2(*a*). In response to the influx at the boundaries, the membrane deforms out of plane as the protein density increases; see figure 2(*b*). Once the flux returns to zero, diffusion homogenizes the protein, and the membrane height begins to decrease again. This effect is observed clearly by looking at the deformation at the center of the string as a function of time in figure 2(*c*), which closely follows the dynamics of the boundary flux show in figure 2(*a*).

**Figure 2:**
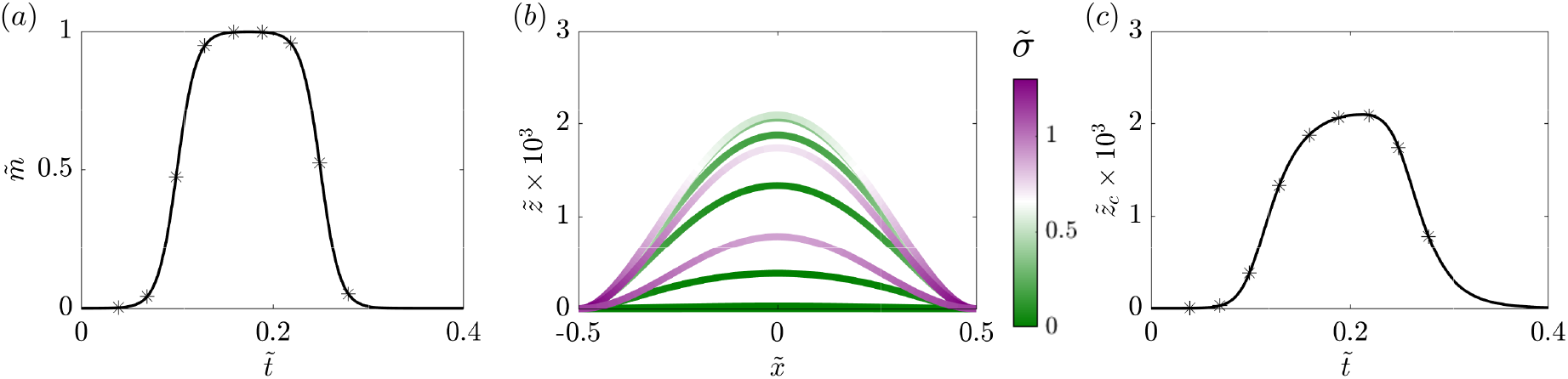
Evolution of membrane deformation and protein distribution when a influx of protein is prescribed at both boundaries. (*a*) Dimensionless boundary protein flux as a function of time. (*b*) Distribution of protein density plotted on the deformed membrane for 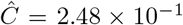. (*c*) Time evolution of the maximum membrane deflection 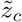 for 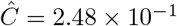. Symbols in panels (a) and (c) correspond to the times shown in (b).

In both examples of figures 1 and 2, we note that the protein distribution becomes uniform at long times (in the absence of any boundary flux), and as a result the membrane returns to its flat reference shape. At first glance, this result seems counter-intuitive since there is a non-zero density of curvature-inducing proteins on the membrane. But as we showed previously, for a uniform distribution of proteins with no-flux boundary conditions on the membranes, minimal surfaces are admissible solutions for the membrane geometry [49, 50]. In this particular case, a flat membrane is the admissible solution for the boundary conditions associated with 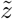.

### 4.2 Two-dimensional simulations

#### 4.2.1 Numerical implementation

We solved the set of governing equations (32)–(35) in two dimensions inside a square domain using a finite-difference technique that we outline here. Our numerical scheme is second order in space and first order in time. We note that time only appears explicitly in the advection-diffusion equation (35) for the protein density: we solve it using a semi-implicit scheme wherein the linear diffusion term is treated implicitly while the nonlinear advective terms and curvature-induced transport terms are treated explicitly. The remaining governing equations are all elliptic in nature and can be recast as a series of Poisson problems as we explain. First, we note that the shape equation (34) is biharmonic and can thus be recast into two nested Poisson problems providing the shape 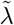 at a particular time step. To solve for the surface tension λ, we take the divergence of the tangential momentum balance (33) and combine it with the continuity equation (32) to obtain the Poisson equation

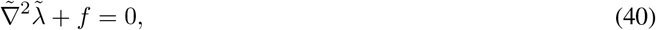

where

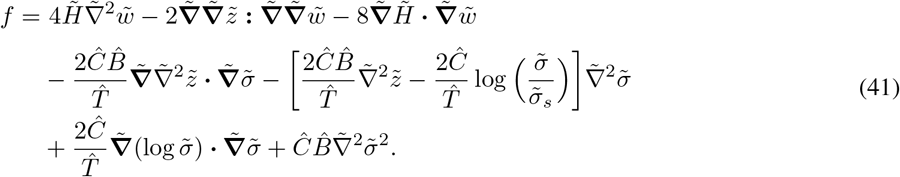

Note that there is no natural boundary condition on 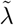 at the edges of the domain. To approximate an infinite membrane, we first estimate the tension along the four edges using the integral representation

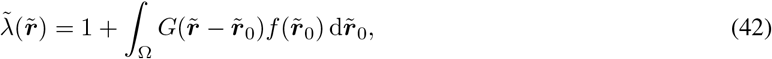

where *G*(***r***) = − log *r*/2*π* is the 2D Green’s function for Poisson’s equation in an infinite domain. The calculated tension along the edges is then used as the boundary condition for equation (40), where the normal velocity component at the current time step *k* is calculated as

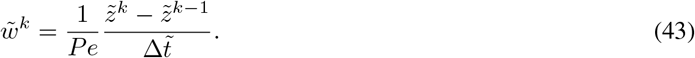

With knowledge of the membrane tension, the tangential momentum balance (33) then provides two modified Poisson problems for the in-plane velocity components. Note that the equations for 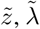 and 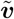 are nonlinearly coupled through the various forcing terms in their respective Poisson problems. To remedy this problem, we iterate their solution until every variable converges with a tolerance limit of 5 × 10^−7^ before proceeding to the next time step. All the results presented below were obtained on a spatial uniform grid of size 201 × 201 and with a dimensionless time step of 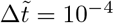. We used Fortran 90 for compiling and running the algorithm.

#### 4.2.2 2D simulation results

Using the numerical scheme described above, we solved the linearized two-dimensional governing equations (32)–(35) for different initial conditions. In all cases, the boundary conditions for the membrane shape were set to 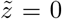 and 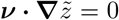 where ***ν*** is the normal to the edge of the domain, and no-flux boundary conditions were enforced on the protein distribution. We considered three different initial conditions for the protein density as depicted in figure 3, namely: a single circular patch at the center of the domain (figure 3(*a*)), two indentical patches placed at diametrically opposite ends of the domain (figure 3(*b*)), and four patches centered in each quadrant of the domain (figure 3(*c*)). The total mass of protein is the same in all three cases, only the initial spatial distribution is different. For the velocity and tension, we maintain open boundary conditions as noted in § 4.2.1.

**Figure 3:**
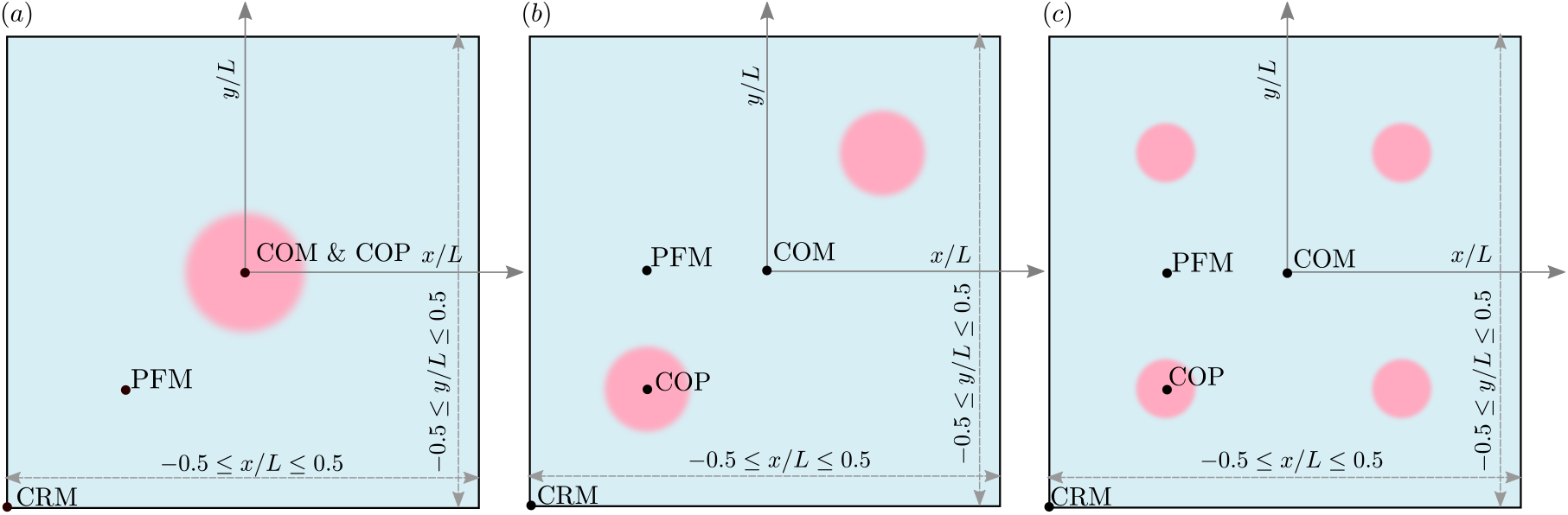
System set up and initial condition used in 2D simulations. All 2D simulations are performed for a linearized Monge patch. We simulate the dynamics for three different initial distributions of proteins as shown. The total area fraction of protein is same for the three cases, with proteins covering 10% of the total area. (*a*) Single patch of protein placed at the center (0, 0)). (*b*) Two patches of protein placed at diametrically opposite positions with center locations (−0.25, −0.25) and (0.25, 0.25). (*c*) Four patches of proteins placed at four diagonal positions: (−0.25, −0.25), (−0.25, 0.25),(0.25, −0.25) and (0.25, 0.25). The following abbreviations are used in subsequent figures to track the system behavior: COM: center of the membrane, COP: center of the patch, CRM: corner of the membrane, and PFM: protein-free membrane.

We tracked the dynamics of the membrane shape, protein distribution, membrane tension, and velocity for a single patch of proteins corresponding to figure 3(*a*) in figure 4. The initial membrane configuration is bent to accommodate the initial distribution of proteins (figure 4(*a*)), and the membrane tension for this initial distribution is heterogeneous as seen in figure 4(*d*), consistent with our previous results [32, 9]. Over time, the proteins diffuse from the center of the patch across the membrane, tending towards a homogeneous distribution (figure 4(*b*, *c*)), and this process is accompanied by a reduction in the membrane deflection. The homogenization of proteins results in homogenization of the membrane tension, which approaches its value at infinity (figure 4(*e, f*)). The tangential velocity is directed outward (figure 4(*g, h, i*)), and the dimensionless magnitude of the maximum velocity in figure 4(*g*) is 4.8 × 10^−3^. Expectedly, the magnitude of this velocity decreases with time as seen in figure 4(*h, i*).

**Figure 4:**
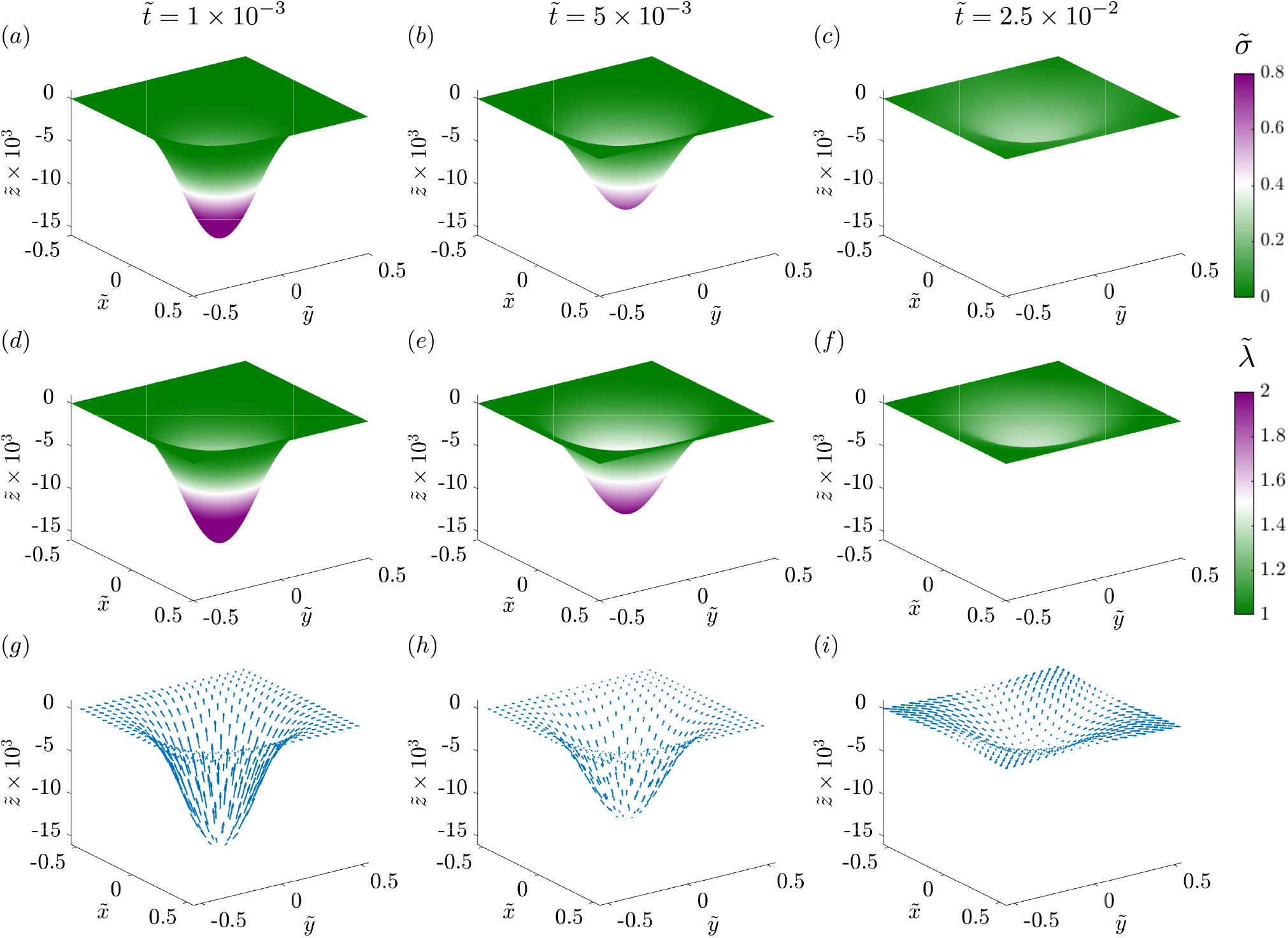
Dynamics of the evolution of membrane shape, protein distribution, membrane tension, and tangential velocity field for a single patch of protein at three different times. Distributions of membrane protein density are shown at dimensionless times 1 × 10^−3^ (*a*), 5 × 10^−3^ (*b*), and 2.5 × 10^−2^(*c*). Distributions of membrane tension at the same non-dimensional times 1 × 10^−3^, (*d*), 5 × 10^−3^(*e*), and 2.5 × 10^−2^(*f*). Tangential velocity fields shown at the same non-dimensional times 1 × 10^−3^ (*g*), 5 × 10^−3^(*h*), and 2.5 × 10^−2^ (*i*). The magnitude of the maximum dimensionless tangential velocity is 2.8 × 10^−3^.

When the proteins are distributed in two and four patches as shown in figure 3(*b, c*), we found that the overall behavior of the system was quite similar to a single patch with some changes to the dynamics. First, because each patch had a lower density of proteins (half or quarter), the initial deformation was smaller and the protein distribution homogenized faster than in the case of a single patch (figure 5(*a, b*)). Similarly, the typical magnitude of membrane tension variations (figure 5(*c, d*)) and of the tangential velocity field (figure 5(*e, f*)) was also smaller to begin with and the system attained the homogeneous distribution rapidly.

**Figure 5:**
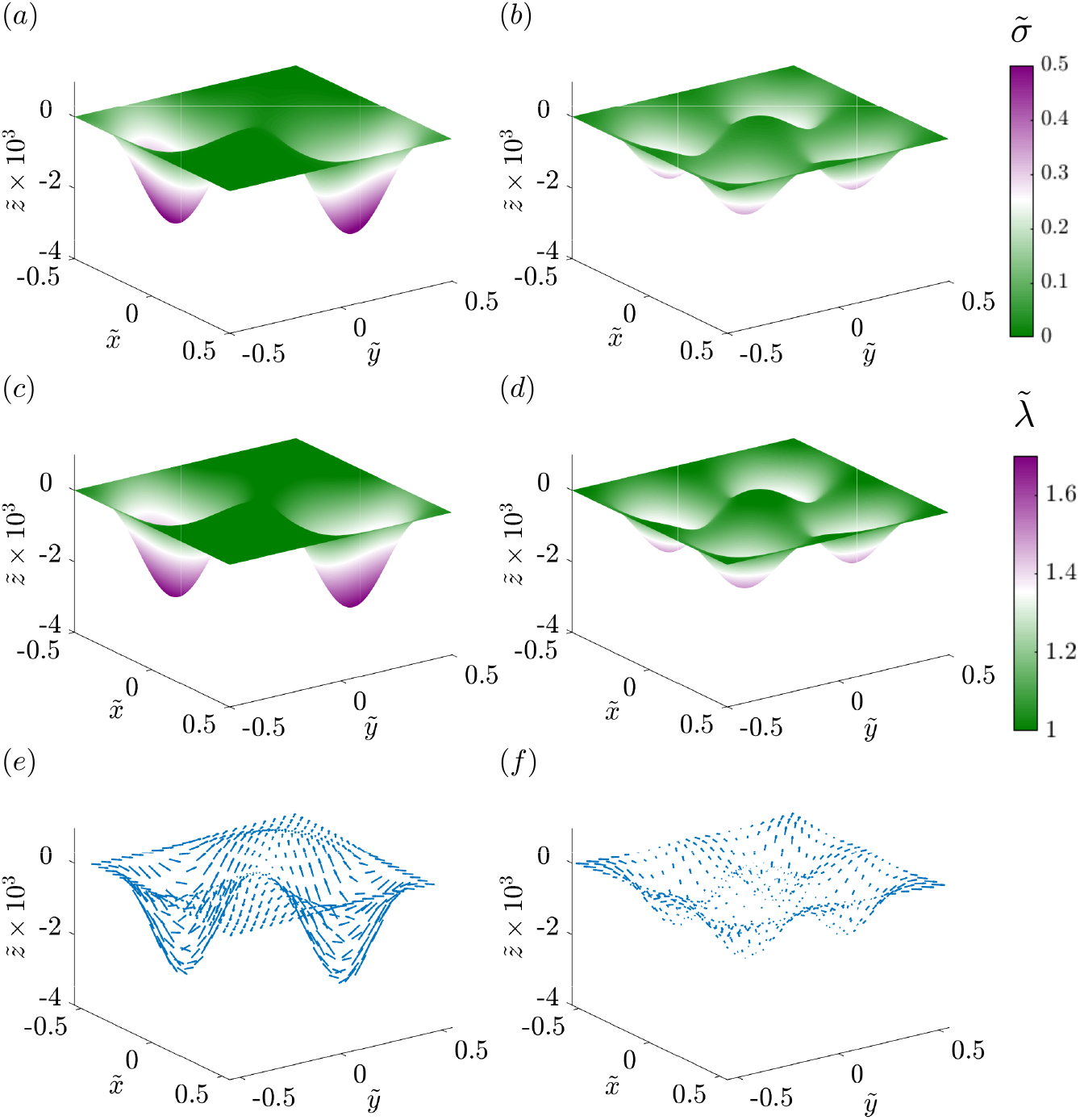
Dynamics of the evolution of membrane shape, protein distribution, membrane tension, and tangential velocity field for two and four patches of protein at dimensionless time 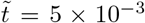. The left column shows the distribution of protein density (*a*), membrane tension (*c*), and tangential velocity (*e*) for two patches of protein. The right column shows the distribution of protein density (*b*), membrane tension (*d*), and tangential velocity (*f*) for four patches of protein. The magnitude of maximum dimensionless tangential velocity is 2.2 × 10^−2^ in the case of two patched, and 4.1 × 10^−3^ in the case of four patches.

To compare the effects of one, two, and four patches directly, we plotted the membrane deformation (figure 6), membrane protein distribution (figure 7), and membrane tension distribution (figure 8) at different locations for each case. The initial deformation is different for the different cases because of the differences in the local density of proteins. For a single patch, we observed that the maximum deformation occurs at the center of the patch (COP) and it takes a longer time for this deformation to go to zero in the case of a single patch compared to multiple patches (compare figure 6(*a*) to figure 6(*b, c*)). in the case of two and four patches, we also observe a small positive deformation at the center of the membrane (COM) and in the protein-free membrane (PFM). This can be explained by the fact that the continuity conditions of the surface will result in a small but upward displacement in protein-free regions in response to the large downward displacement in the regions where the proteins are initially present.

**Figure 6:**
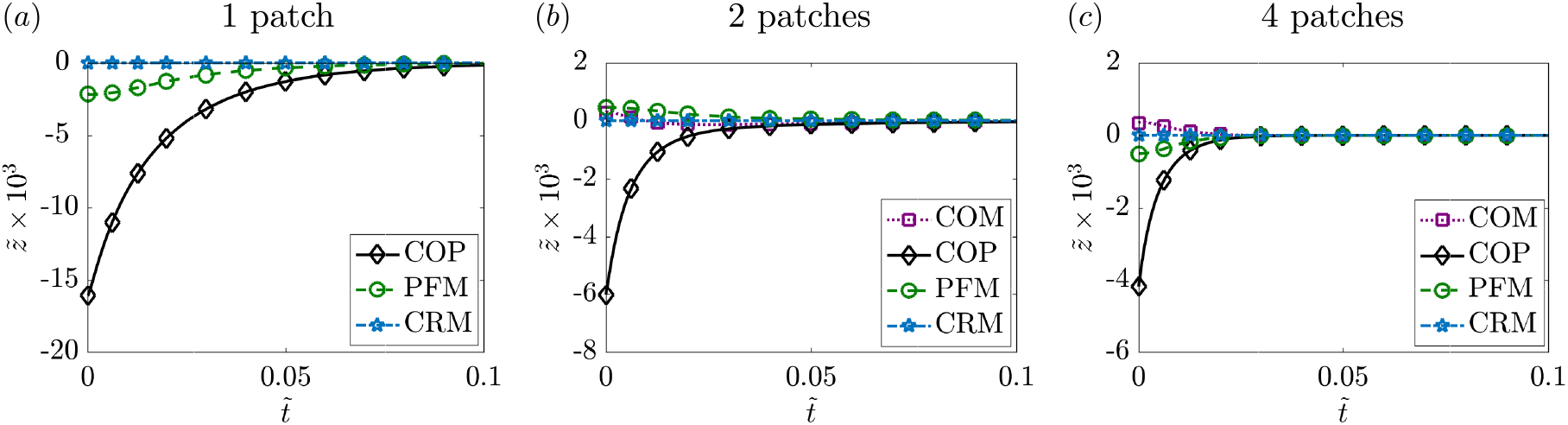
Temporal evolution of the membrane deflection at the various locations defined in figure 3 for a single patch of protein (*a*), two patches (*b*), and four patches (*c*).

**Figure 7:**
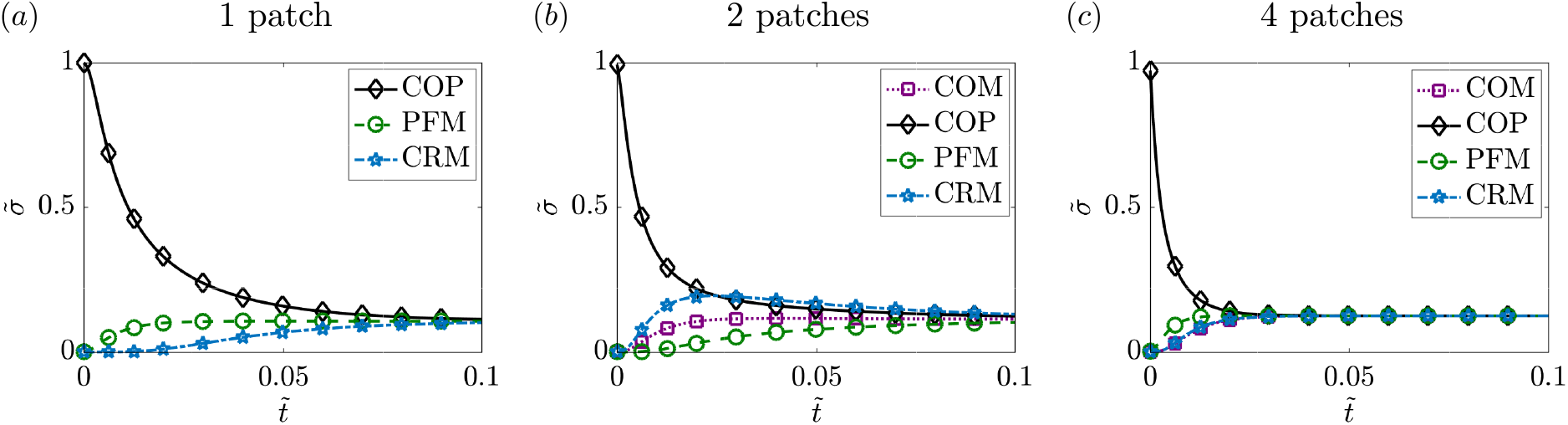
Temporal evolution of the protein density at the locations defined in figure 3 for a single patch of protein (*a*), two patches (*b*), and four patches (*c*).

**Figure 8:**
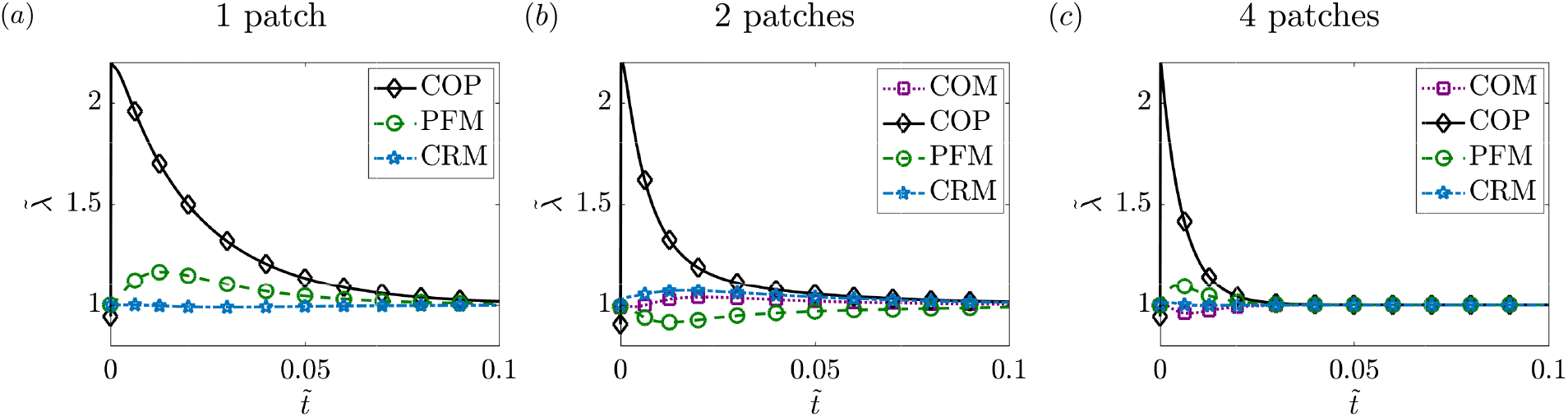
Temporal evolution of the membrane tension at the locations defined in figure 3 for a single patch of protein (*a*), two patches (*b*), and four patches (*c*).

Comparing the protein dynamics for one, two, and four patches, we observed that increasing the number of patches decreases the time it takes for the protein distribution to homogenize across the membrane domain (figure 7). Thus, although membrane bending and protein distribution are coupled, the distribution of multiple patches weakens the coupling and promotes rapid homogenization of the membrane proteins. While the steady state protein distribution is the same in all cases, the dynamics with which the protein-free regions show an appreciable increase in proteins also depends on the initial distribution of proteins. For example, in the case of a single patch, CRM takes much longer to reach steady state compared to the case of two or four patches. Figure 7 shows that for the initial conditions of a single patch and of four patches of protein, 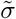 approaches the uniform protein density monotonically. But for the case of two patches, the time evolution of the protein density is not monotonic, and we found the density of protein at the corner of the membrane (CRM) exceeds the density at the center of the patch (COP) for a brief time interval. We investigated this phenomenon further and studied the dependence on intra-membrane flow by varying the Péclet number in § 4.2.3.

Similar dynamics are observed for the membrane tension as well. Figure 8 shows that the membrane tension takes a larger time to reach its steady value for the case of one or two patches when compared to four patches (compare figures 8(*a, b, c*)). The initial rise in the membrane tension corresponds to the inviscid response of the membrane to the curvature-inducing protein distribution, while from the next time step onwards tension changes primarily due to viscous effects.

#### 4.2.3 Effect of Péclet number on membrane dynamics

Next, we investigated the effect of fluid advection in the case of two patches by varying the Péclet number *Pe* in figure 9. We observed that at the center of the patch (figure 9(*a*)) there was no observable effect of varying the Péclet numbers on the temporal evolution of the protein density. However, at the center of the membrane (figure 9(*b*)) and in the protein-free membrane (figure 9(*c*)), we observed that increasing Péclet number had a small effect on the dynamics of the protein density, particularly at long times.

**Figure 9:**
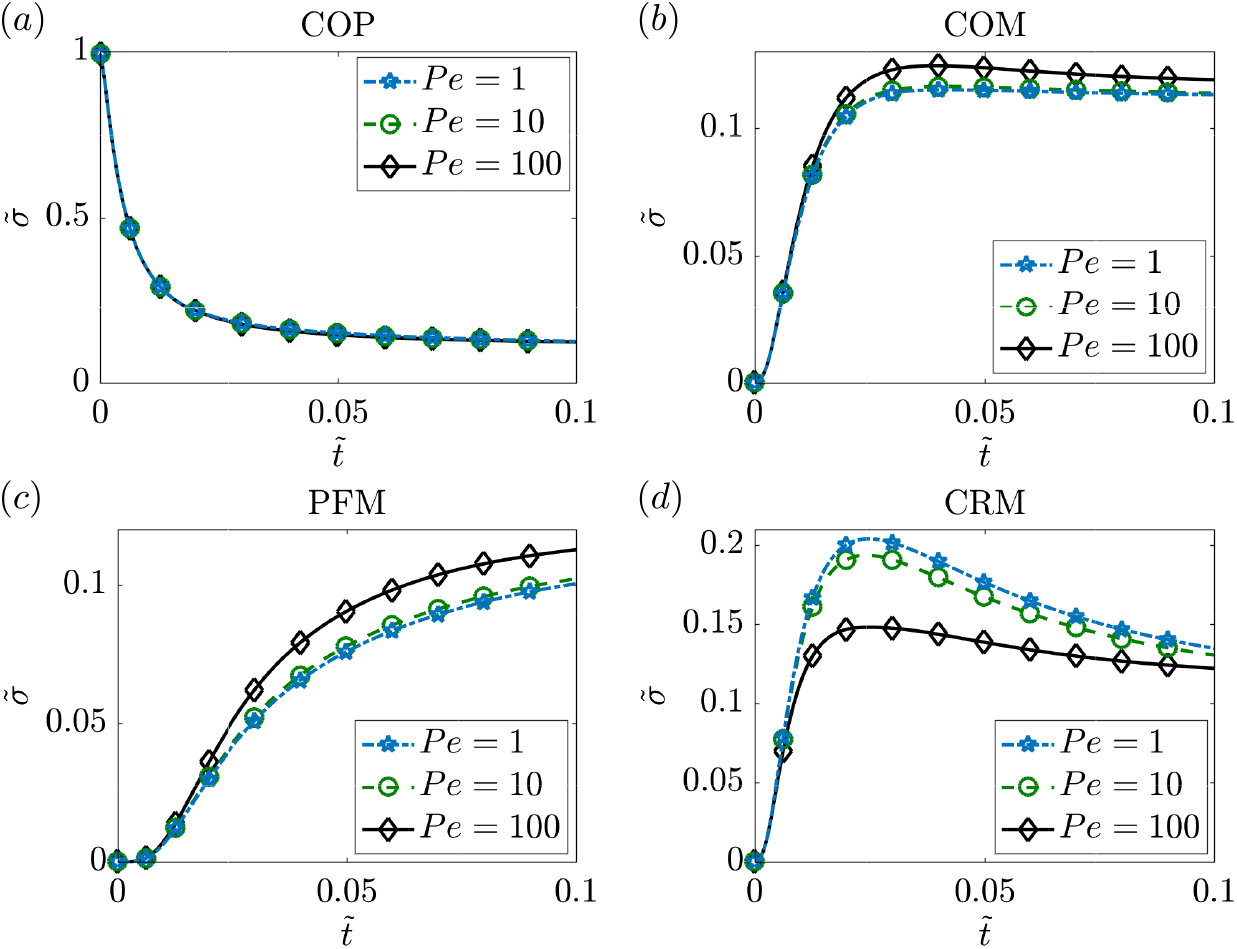
Temporal evolution of local protein density for different values of the Péclet number in the case of two patches of protein. The protein density is measured at the four locations defined in figure Figure 3: COM (*a*), COP (*b*), PFM (*c*) and CRM (*d*).

The effect of increasing the Péclet number was most dramatic at the corner of the membrane (Figure 9(*d*)), where the initial rise in the protein density was found to be similar for all three values of *Pe*, but the increase resulted in a higher value for lower *Pe*. Eventually, 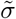 at the corner decreases towards the mean value of 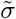 over time. Thus, the coupling between lipid flow and protein diffusion seems to have a larger impact on transport in the regions that are initially protein-free.

To further investigate the role of convective transport, we tracked the separation distance 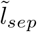 between the centers of mass of two effective patches (*l_sep_*) as a function of time in figure 10. The center of mass of a patch is formally defined as

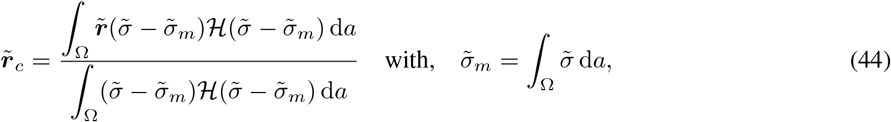

where the effective extent of the patch is defined using the Heaviside function 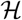 as the area where protein density exceeds its mean value. We observed that 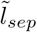 increases with time and decreases with increasing Péclet number (figure 10). This can be explained from the velocity profile for two patches in figure 5(*e*). The direction of the velocity is towards the center of the membrane in the area where the patch is located. Therefore, the advective transport due to the lipid tends to reduces the separation otherwise caused by diffusion. Since, with increasing *Pe* the effect of flow increases, the effect of separation decreases for higher values Péclet number as shown in figure 10. This also explains the decrease of density of the protein at the corner of the membrane (CRM) and the increase of the protein density at the initial protein-free area (PFM) and center of the membrane (COM) with higher value of *Pe* as found in figure 9.

**Figure 10:**
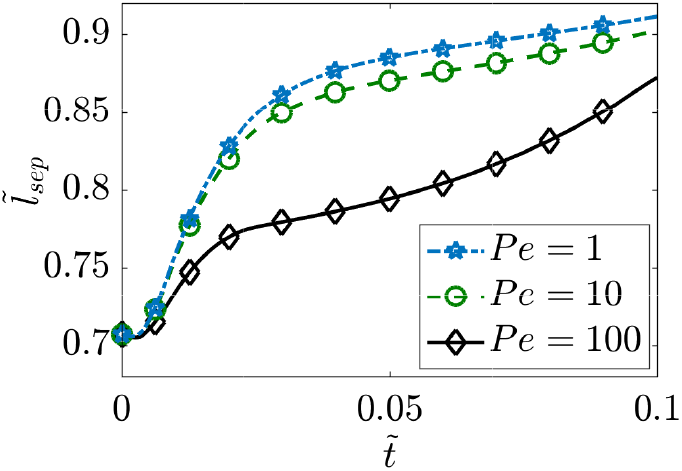
Evolution of the separation distance between the centroids of the protein patches in the case of two patched and for three different values of the Péclet number.

We can further understand the dynamics of the separation distance between the two patches by considering the diffusion of a protein patch in one triangular half domain of lipid. This triangle is bounded by two of the domain boundaries and by the diagonal of the square domain that passes in between the two patches. The diagonal line is also a line of symmetry, and thus behaves as an effective no-flux boundary for the triangular half of the domain. Therefore, each triangular half-domain is subject to the no-flux condition on its three sides. In this half domain, the semicircular half patch of protein facing the corner of the membrane (CRM) diffuses to a smaller area compared to the other semicircle that faces the center of the membrane (COM). This results in an effective larger protein gradient towards CRM. Therefore, the protein density shifts towards the protein-free corner and results in an effective shift of the patches towards the corners of the membrane.

## 5 Conclusions and discussion

In this work, we have derived and analyzed the governing equations for the protein-induced deformation of a lipid membrane coupled with the protein diffusion and in-plane viscous flow of the lipids. The coupling between diffusion and lipid flow completes the description of the key transport phenomena involved in lipid membranes. We conducted simulations in 1D and 2D and further quantified the relationship between membrane bending and protein diffusion. The major conclusions from our study are that lipid flow and membrane protein diffusion, when coupled, can alter the dynamics of membrane protein distribution at different locations. We find that as the protein diffuses from an initial locally concentrated patch, the membrane deformation decreases and these dynamics are also related to the diffusion coefficient of proteins on the membrane. The flow of lipids also seems to induce a separation dynamics that depends on the Péclet number of the system when multiple patches are present.

Previously, we elaborated on the need for coupling between the viscous and elastic effects for the calculation of the Lagranage multiplier associated with the incompressibility constraint of the membrane [32, 29]. Here, we build on that framework to include protein diffusion. The coupled interaction between elasticity, diffusion, and viscous flow now fully describes the equations associated with the Lagrange multiplier λ, reinforcing its interpretation as a surface pressure [42, 29].

There have been many studies focused on modeling membrane-protein interactions [51, 12]. Here, we show that coupling the viscous flow of lipids on the membrane is important for modulating the dynamics of the system and fully describing interfacial transport phenomena. Future efforts will focus on adsorption of proteins from the bulk [52, 53] and phase separation of proteins to identify the coupling between lipid flow and chemical energies associated with these processes on an elastic membrane. Such theoretical developments not only have implications for our understanding of biological membranes, but also have the potential to impact curvature-driven, directed assembly in colloids and liquid crystals suspended in fluids, and particle interactions at interfaces between immiscible fluids and soft materials, enabling directed design and engineering of the next-generation of reconfigurable systems in soft matter [54, 55].

## Acknowledgements

The authors would like to thank Prof. David Steigmann for initial discussions on the model development. This work was supported in part by ONR N00014-17-1-2628 to P.R.

